# Origin, evolution and diversification of plant mechanosensitive channel of small conductance-like (MSL) proteins

**DOI:** 10.1101/2022.12.09.519821

**Authors:** Zaibao Zhang, Fan Ye, Tao Xiong, Jiahui Chen, Jiajia Cao, Yurui Chen, Tian Luo, Sushuang Liu

**Affiliations:** School of Life and Health Science, Huzhou College, Huzhou, Zhejiang, China; College of International Education, Xinyang Normal University, Xinyang, Henan, China; College of Life Science, Xinyang Normal University, Xinyang, Henan, China

**Keywords:** mechanosensitive ion channel, MscS-like (MSL), molecular evolution, origin, expansion

## Abstract

Mechanosensitive (MS) ion channels provide efficient molecular mechanism for transducing mechanical forces into intracellular ion fluxes in all kingdoms of life. The mechanosensitive channel of small conductance (MscS) was one of the best-studied MS channels and its homologs (MSL, MscS-like) were widely distributed in cell-walled organisms. However, the origin, evolution and expansion of MSL proteins in plants are still not clear. Here, we identified more than 2100 MSL proteins from 176 plants and conducted a broad-scale phylogenetic analysis. The phylogenetic tree showed that plant MSL proteins were divided into three groups (I, II and III) prior to the emergence of chlorophytae algae, which was consistent with their specific subcellular localization. MSL proteins were distributed unevenly into each of plant species, and four parallel expansion was identified in angiosperms. In Brassicaceae, most MSL duplicates were derived by whole-genome duplication (WGD)/segmental duplications. Finally, a hypothetical evolutionary model of MSL proteins in plants was proposed based on phylogeny. Our studies illustrate the evolutionary history of the MSL proteins and provide a guide for future functional diversity analyses of these proteins in plants.

## Introduction

All living organisms are subjected to various external and internal mechanical stresses, including gravity, touch, sound and osmotic shock. How mechanical forces are sensed by cells is a long-standing question in biology. One of the most universal mechanisms for cells to respond to mechanical stimuli is the use of mechanosensitive (MS) ion channels (Cox et al. 2015). MS channels are transmembrane proteins that exist in all kingdoms of life. The primary function of MS channels is to provide a conductive pore in response to mechanical stimulation, allowing ions to flow across the membrane down their electrochemical gradient (Anishkin and Kung 2005).

Many types of MS ion channels have been identified in different organisms, including mechanosensitive ion channel of small conductance (MscS) (Malcolm and Maurer 2012; Pivetti et al. 2003), mechanosensitive ion channel of large conductance (MscL) (Martinac 2001), two pore potassium (TPK) (Honoré 2007), Mid1-complementing activity (MCA) (Kurusu et al. 2012), and piezo (Volkers et al. 2015). Different MS channels displayed highly divergent in conductance, ion selectivity, and/or sensitivity to the direction of activation pressure. MscS is a nonselective stretch-activated channel which is gated by membrane tension (Booth et al. 2011; Kung et al. 2010). *Escherichia coli* has six MscS paralogs: archetypal MscS (yggB), potassium-dependent MscK (kefA), MscM (YjeP), YbdG, YbiO and YnaI (Schumann et al. 2010; Edwards et al. 2012). These bacteria MscS channels have different activation thresholds and channel conductance, protecting cells from osmotic stress by providing a conduit for the release of osmolytes from the bacterium (Cox et al. 2018; Levina et al. 1999; Boer et al. 2011).

MscS family members are highly divergent in their topology and domain structure. The crystal structure of *E. coli* archetypal MscS (PDB: 2OAU – EcMscS, PDB) was resolved (Bass et al. 2002), and it was characterized by three N-terminal transmembrane (TM) helices followed by a large hydrophilic cytoplasmic domain. The key feature of the EcMscS structure is the pore lining TM helix, TM3, which forms a hydrophobic channel pore and shows the highest homology in MscS-like channels (Pivetti et al. 2003; Balleza and Gómez-Lagunas 2009). TM3 comprises two regions, TM3a and TM3b, which are separated by a distinctive kink at residue G113. A comparison of the open-state versus closed-state structures of EcMscS showed that gating involves swinging a tension-sensitive paddle made up of the TM1/TM2 helices and twisting TM3a about G113 (Nomura et al. 2006; Malcolm et al. 2011).

MscS homologs are widely dispersed in bacterial, archaeal, fungal and plant genomes, but not in animal genomes. In Arabidopsis, ten MscS homologs were identified, named as MscS-like proteins (MSLs) (Haswell 2007). The Arabidopsis MSLs were divided into three phylogenetic groups, consist with their different subcellular localization and topology (Hamilton et al. 2015b). Group I (AtMSL1) and group II MSLs (AtMSL2 and AtMSL3) were localized in the inner membrane of mitochondria and chloroplast, respectively (Haswell 2007; Haswell and Meyerowitz 2006; Haswell et al. 2008). Group III MSLs (AtMSL4-10) were localized in the plasma or endoplasmic reticulum (ER) membrane (Haswell 2007; Hamilton et al. 2015b; Guerringue et al. 2018). Both group I and group II MSLs contained five TM helices, while group III MSLs contained six TM helices. The distinct cellular localizations implicate diverse physiological functions of plant MSLs. Loss of *AtMSL1* increased mitochondrial oxidation under abiotic stresses, indicating that AtMSL1 is crucial for regulating mitochondrial redox status under abiotic stress (Lee et al. 2016). AtMSL2 and AtMSL3 colocalize with the plastid division protein AtMinE and function redundantly in maintaining plastid shape, size and division (Haswell and Meyerowitz 2006; Veley et al. 2012; Wilson et al. 2011). *AtMSL8* is specially expressed in pollen, and plays essential roles in pollen hydration and pollen tube growth during fertilization (Hamilton and Haswell 2017; Hamilton et al. 2015a). Both loss of function and overexpression of *AtMSL8* lead to reduced pollen germination and low fertility. *AtMSL10* is expressed in root and form a heteromeric channel with AtMSL9 (Haswell et al. 2008; Peyronnet et al. 2008). Both loss of function and overexpression of *AtMSL10* lead to growth retardation and ectopic cell death (Basu et al. 2020). AtMSL10 is also involved in the wound-triggered early signal transduction and plays a positive regulatory role in biosynthesis of jasmonic acid (Zou et al. 2016).

The MscS homologs were discovered in Bacteria, suggesting that MscS existed in the early stages of evolution. However, its identification and functional analyses in plants are still limited. In order to explore the origin, characterization and diversification of plant MSLs, we sought to build a comprehensive phylogeny of plant MSLs. 2123 MSL proteins were identified from 176 plants. Based on the phylogenetic tree, we explored the origin and divergence of the plant MSL proteins. Three MSL groups (I, II and III) were identified in plants, and the divergence of these three MSL groups can be traced back to chlorophytae algae. In addition, a wide phylogenetic architecture of angiosperm MSLs were constructed, and the MSLs in angiosperms were further classified to 4 clades: MSL1, MSL2/3, MSL4-8 and MLS9/10. Finally, we discussed the possible evolutionary relationships of the MSL proteins in plants.

## Results and discussion

### MSL proteins were identified in all lineages of plant

For constructing a comprehensively phylogeny of MSL proteins in plants, we used the Hidden Markov Models (HMM) algorithm and BLASTP search to identify MSL proteins in genome-sequenced species, including 2 chlorophytic algae, 6 charophytes, 5 bryophytes, 2 ferns, 5 gymnosperms and 156 angiosperms (Supplementary Table 1). The retrieved proteins were checked with SMART and PFAM, and the candidates containing the MS_channel domain were regarded as ‘true’ MSLs. In total, 2123 MSL proteins were retrieved from 176 plant species (Table 1). The copy number of MSL proteins varies among different plant lineages, ranging on average from 5.5 copies in chlorophyta, 4.7 in bryophytes, 9.0 in ferns, 6.0 in gymnosperms, to 12.7 angiosperms (Table 1). These data suggest that the MSL proteins were expanded in angiosperms. Among the angiosperms examined, more MSL proteins were identified in dicots than that in monocots, with an average of 9.2 in Poaceae, 12.5 in Brassicaceae, 12.0 in Leguminosae and 11.1 in Rosaceae (Table 1). These results indicate that MSL proteins were expanded in dicots.

**Table 1.** The number of MSL proteins in green plants.

### Evolutionary origin of MSL proteins in green plants

MSL proteins were wildly distributed in bacterial, archaeal and plant genomes, but not in animal genomes. The full length of MSLs exhibit little similarity, however, the hydropholic pore-lining TM3 helix and upper cytoplasmic domain display high similarity. Using the pore-lining helix and the conserved cytoplasmic regions of MSLs, a predicted evolutionary tree was constructed among representative members of MSL homologs from bacteria, fungi, protozoa and plants (Figure S1). The plant MSL proteins are separated from bacteria, protozoa and fungi MSLs, and fall into three distinct phylogenetic groups. These results indicated that the divergence of plant MSLs occurred after the emergence of plants. To further explore the evolutionary relationships of MSL proteins in green plants, we reconstructed phylogenetic tree with MSL homologs from 16 represent plant species including green algae *Chlamydomonas reinhardtii*, moss *Physcomitrella patens*, fern *Selaginella moellendorffii*, gymnosperm *Pinus lambertiana*, basal angiosperm *Amborella trichopoda*, monocots (*Oryza sativa, Zea mays*), and dicots (*Arabidopsis thaliana, Arabidopsis lyrata, Theobroma cacao, Glycine max, Camellia sinensis, Coffea canephora*) (Figure 1). The topology of the phylogenetic tree clearly divided plant MSLs into three clades (Clade I, II, III) (Figure 1a), consisting with the three distinct subcellular localizations predictions (Figure 1b). Each clade contains genes from several major lineages of green plants, including algae, mosses, and gymnosperms, indicating that plant MSLs originated in the ancestors of green plants. The angiosperm clade I (MSL1) is monophyletic, indicating that the emergence of MSL1 occurred in the common ancestor of angiosperms. In clade II (MSL2/3), the divergence of MSL2 and MSL3 was observed in dicots, suggesting that the diversification into MSL2 and MSL3 occurred in the ancestor of dicots. More MSL homologs were identified in clade III (MSL4-10), and the divergence of MSL4-8 and MSL9/10 was observed in seed plants (gymnosperms and angiosperms), suggesting that the diversification into MSL4-8 and MSL9/10 occurred in the ancestor of seed plants.

**Figure 1.**
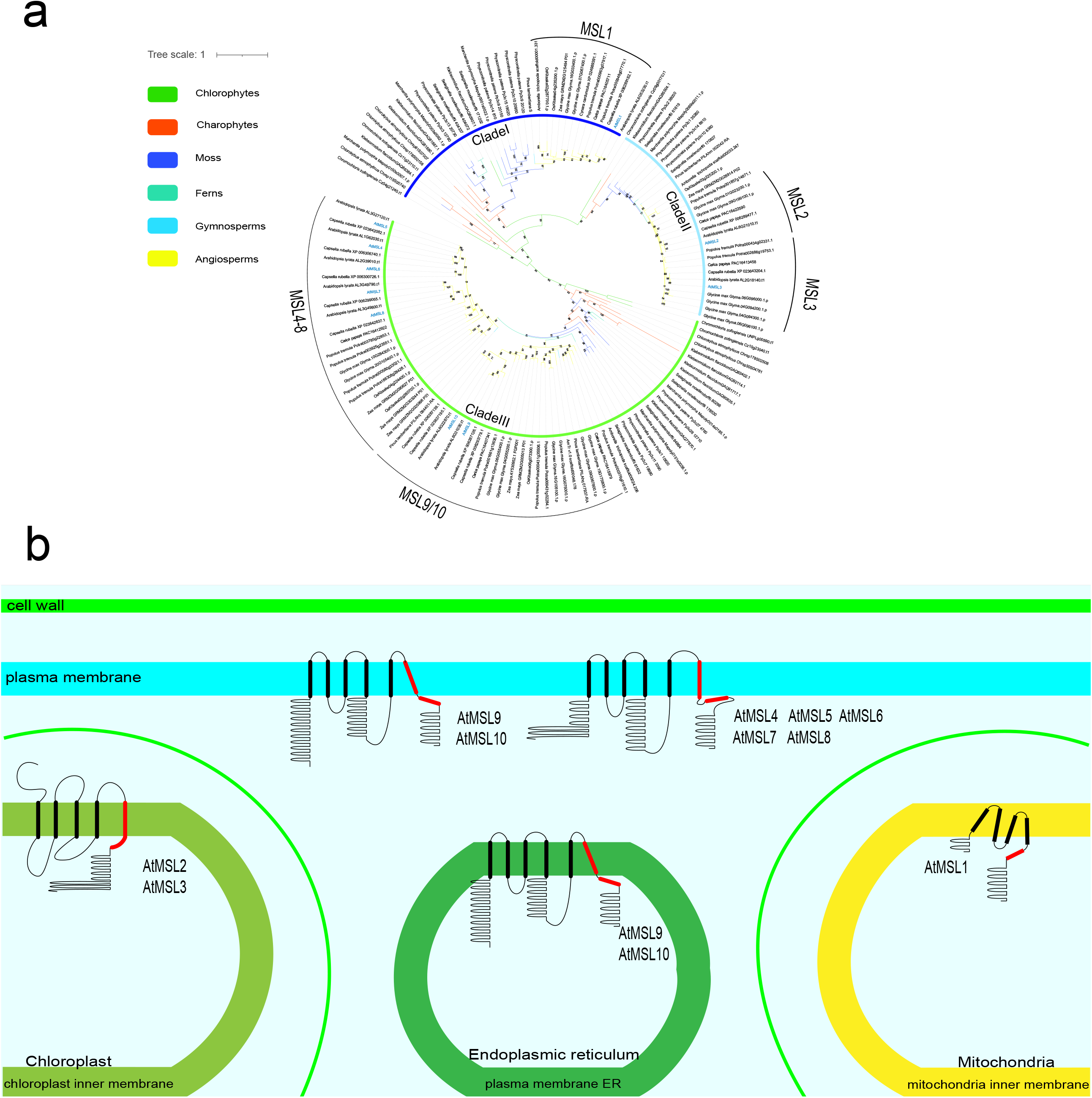
Phylogenetic relationships and subcellular localizations of MSL proteins. (a) An phylogenetic tree of MSL proteins using Bayesian method in green plants. 187 MSL proteins from 3 chlorophytes, 13 charophytes, 5 bryophytes, 1 lycophyte, 2 ferns, 5 gymnosperms and 4 angiosperms were included in phylogeny. (b) Predicted subcellular localization and topology of MSL proteins from *Arabidopsis thaliana* (modify from Hamilton et al., 2015 and Li et al., 2020). Topologies were drawn according to predictions on WoLF PSORT (https://wolfpsort.hgc.jp). The regions of highest homology to *E*.*Coli* MscS TM3 were shown in red.

### Phylogenetic classification of the MSL proteins in angiosperms

To further explore the phylogenetic relationship of MSL proteins in angiosperms, we reconstructed a wide phylogenetic tree with 1971 MSL proteins identified from 155 angiosperm species (Figure 2). The phylogenetic tree shows that angiosperm MSLs were divided into four major groups (MSL1, MSL2/3, MSL4-8, MSL9/10) (Figure 2). MSL1 clade is monophyletic in angiosperm, and many species-specific amplifications of MSL1 were identified, with 6, 5, 7, 13, 7 MSL homologs were identified in rice, *Brachypodium distachyon, Phoenix dactylifera, Dendrobium catenatum, Nicotiana tabacum*, respectively (Figure 2). MSL2/3 was divided into MSL2 and MLS3 in dicots, and most plants have more MSL homologs corresponding to Arabidopsis MSL2 and MLS3 (Figures 2, 3). MSL4-8 and MSL9/10 are lineage-specific paralogs within Brassicaceae (Figures 2, 4, 5). In addition, compared with monocots, more MSL homologs were identified in dicots, indicating that MSLs were expanded in dicot. The topological structure of the gene tree is similar to that of species tree, indicating that these four MSL clades originated independently.

**Figure 2.**
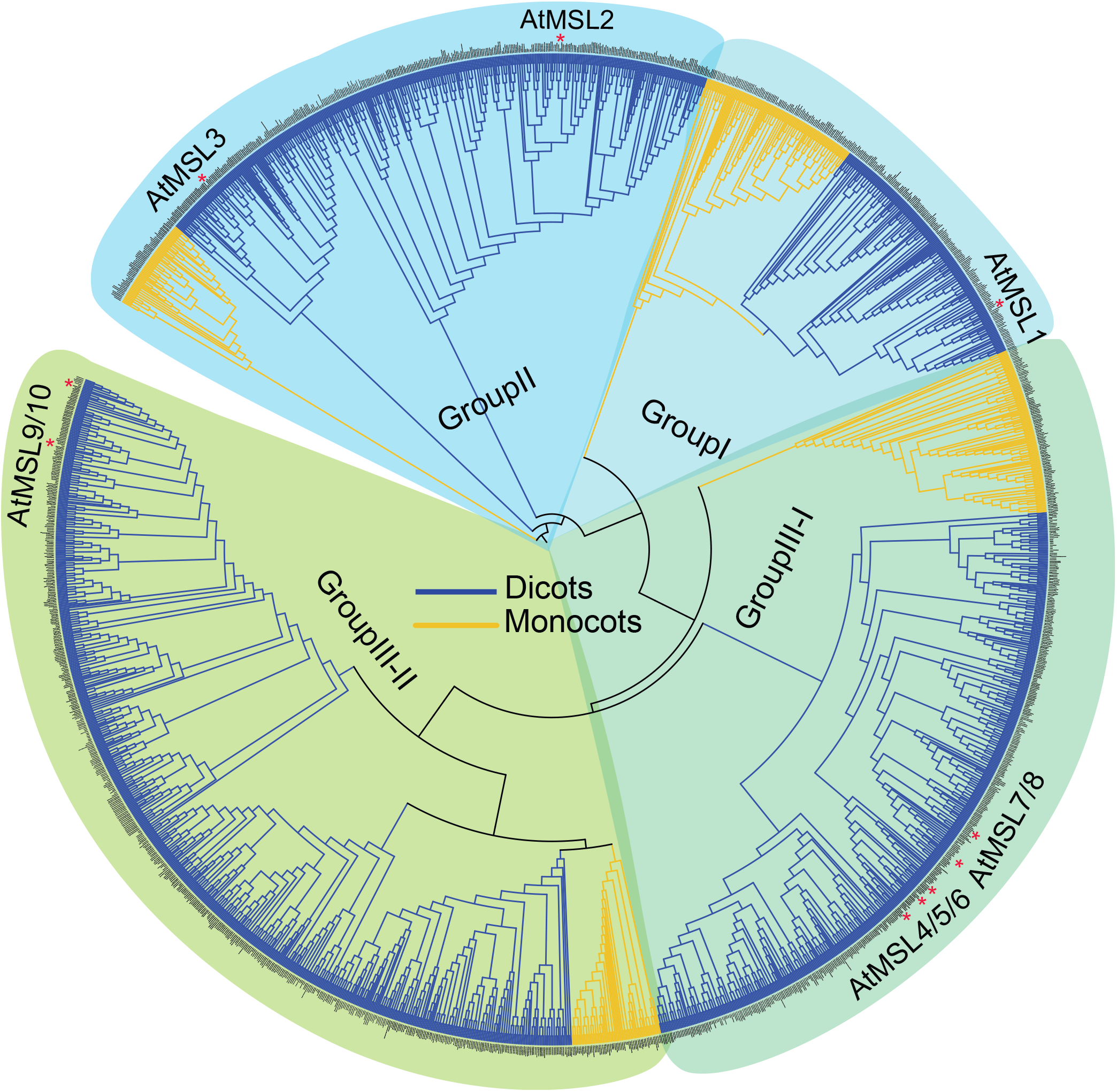
Phylogenetic classification of MSL proteins in angiosperms. The topology shows that MSLs in angiosperms can be classified into 4 sub-groups: MSL1, MSL2/3, MSL4-8, and MSL9/10. Dicots are marked in blue and monocots are marked in yellow.

**Figure 3.**
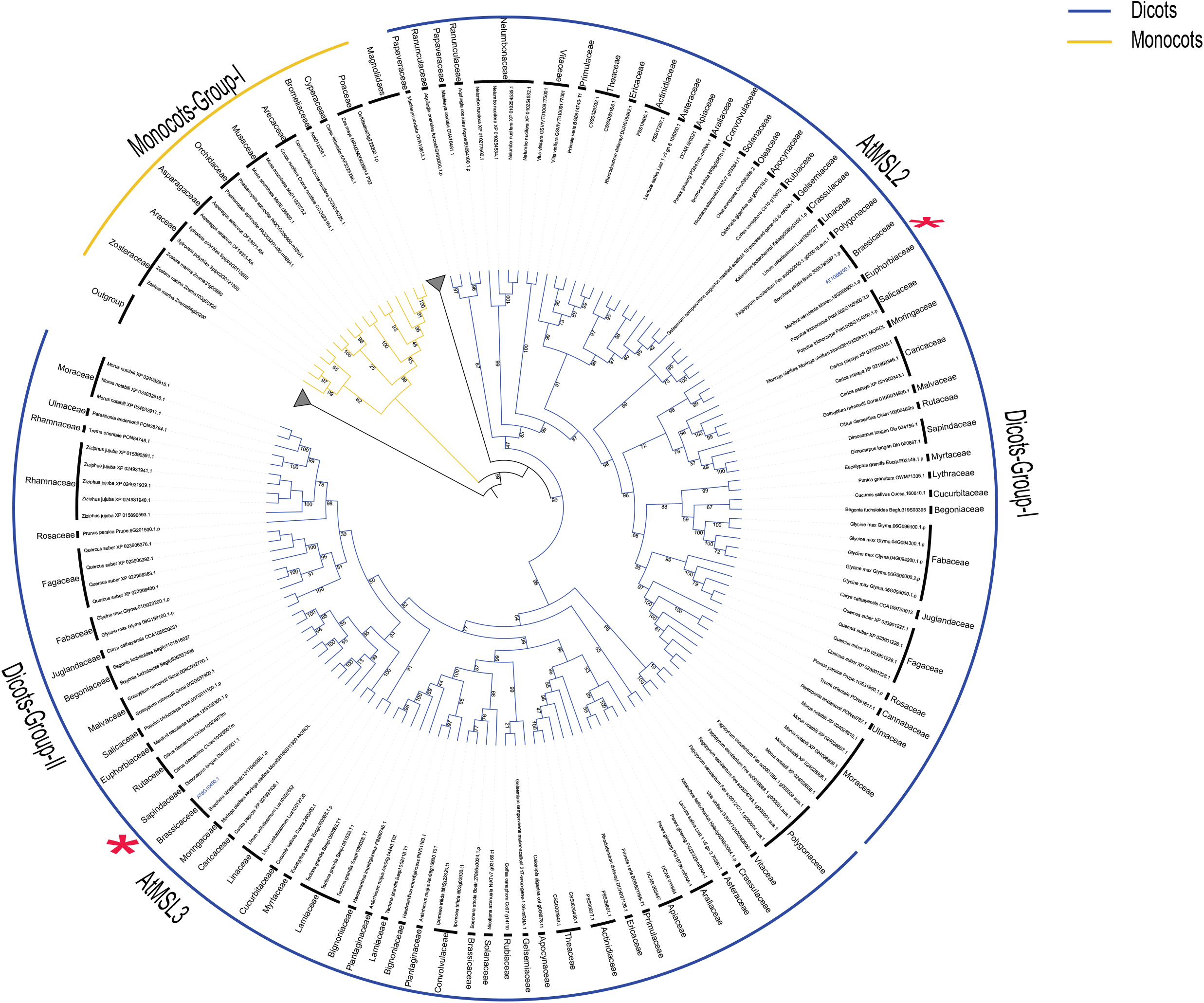
A brief phylogenetic tree showing the MSL2/3 clade in angiosperms. Only selected species were included to represent each order. The topology shows that MSL2/3 in dicots can be clearly classified into two clades, Dicot-Group-I and Dicot-Group-II. Monocots, yellow; Dicots, blue. The outgroup and magnoliids collapsed into a grey triangle.

**Figure 4.**
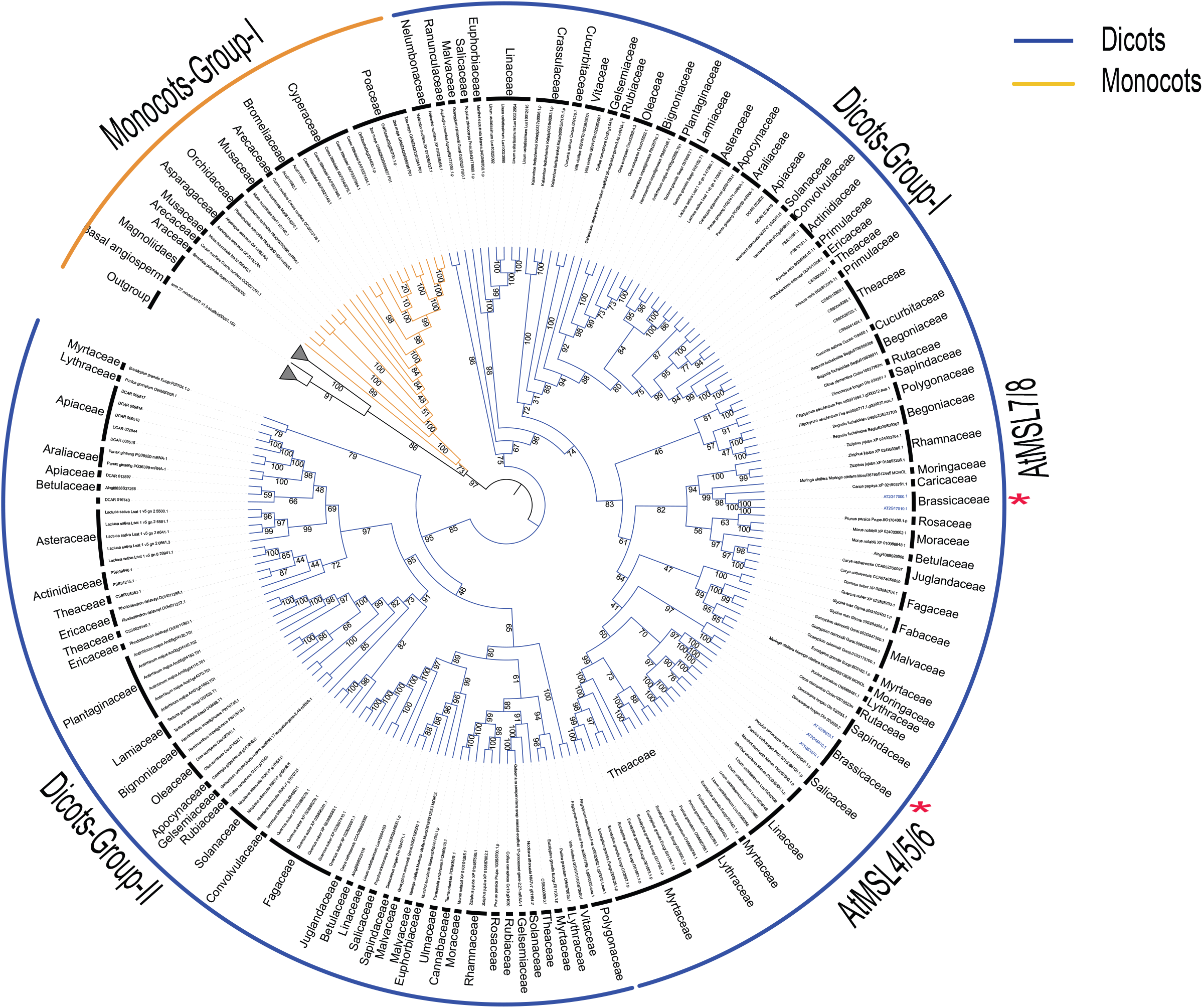
Phylogenetic relationship within the MSL4-8 clade. Only selected species were included to represent each order. The topology shows that MSL4-8 in dicots can be clearly classified into two clades, Dicot-Group-I and Dicot-Group-II. Monocots, yellow; Dicots, blue. The outgroup and magnoliids collapsed into a grey triangle.

**Figure 5.**
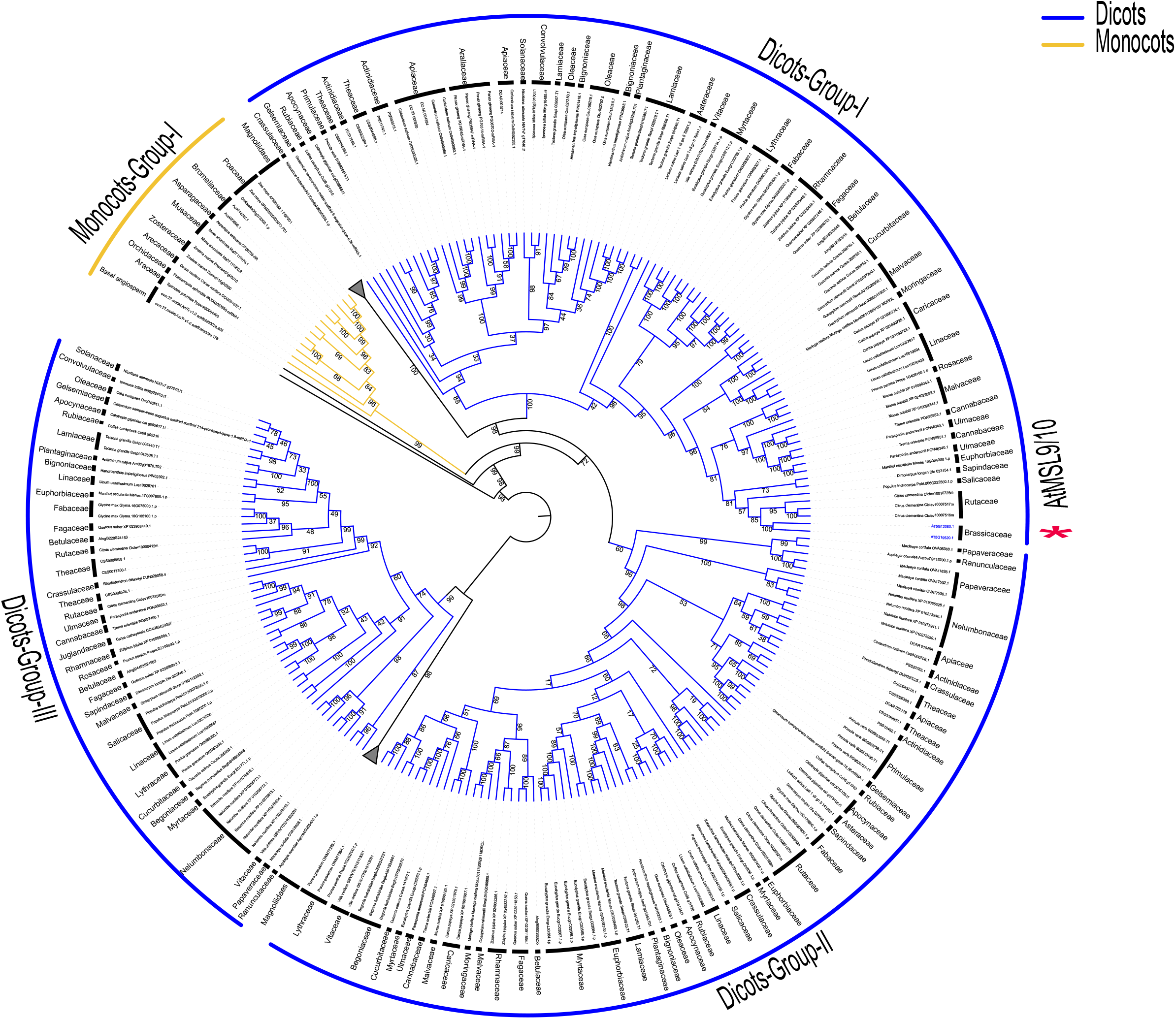
Phylogenetic relationship within the MSL9/10 clade. Only selected species were included to represent each order. The topology shows that MSL9/10 in dicots can be clearly classified into three clades, Dicot-Group-I, Dicot-Group-II and Dicot-Group-III. Monocots, yellow; Dicots, blue. The outgroup and magnoliids collapsed into a grey triangle.

In the MSL2/3 clade, MSL2 and MSL3 shared common dicot lineages and no monocot and magnoliidae species were included, indicating that the divergence between MSL2 and MSL3 occurred before the emergence of dicots and after the monocots/magnoliidae/dicots divergence (Figure 3). In the MSL4/5/6/7/8 (MSL4-8) clade, two sub-branches were identified in dicots: Dicots-Group I and Dicots-Group II (Figure 4). MSL4/5/6/7/8 of all Brassicaceae plants were clustered in Dicots-Group I, and Brassicales MSLs were not appear in Dicots-Group II. In addition, MSL4-8 in Brassicaceae experienced at least three expansion events, resulting in 5 copies of MSL in each species (Figure 4). In the MSL9/10 clade, three sub-branches were identified in dicots: Dicots-Group I, Dicots-Group II and Dicots-Group III (Figure 5). MSL9/10 of all Brassicaceae plants were clustered in Dicots-Group I, and Brassicaceae MSLs were lost in Dicots-Group II and III. The differentiation between MSL9 and MSL10 is due to Brassicaceae-specific duplication (Figure 7).

### Multiple sequence alignment and protein structure prediction

MSL proteins are characterized by a varying number of N-terminal transmembrane (TM) helices followed by a large hydrophilic cytoplasmic domain comprised primarily of ß-sheets (Figure 1) (Bass et al. 2002; Li et al. 2020). The key feature of the MSL proteins is the pore-lining TM helix (TM3 in EcMscS, TM5 in AtMSLs), which is broken into two parts TM3a (TM5a) and TM3b (TM5b). TM5a of AtMSL1 is rich in glycine and alanine residues, which is conserved in group I MSLs and similar to EcMscS (Figure 6) (Li et al. 2020). However, multiple phenylalanine residues are rich in TM5a of group III MSLs, which is different to MSLs of group I and group II (Maksaev et al. 2018). G113 is the kink region of EcMscS and is essential for channel conductance (Bass et al. 2002). However, differential polar residues were identified in the kink region of plant MSLs (Figure 6). R326 and D327 in MSL1 group, R280 and E281 in MSL2/3 group, and G556 and N557 in MSL4-10 group, were identified and proved to be essential for channel conductance (Li et al. 2020; Maksaev et al. 2018; Jensen and Haswell 2012; Schlegel and Haswell 2020). In addition, many group-specific residues were identified and proved to function in MSL molecular and biological role (Figure 6). The A324 and L329 residues were conserved in MSL1 group and worked like a switch for gating and closing of the channel (Li et al. 2020). F553 and I554 residues in AtMSL10 were essential for channel conductance and the stability of the open state of the channel (Maksaev et al. 2018). V273 and L277 of AtMSL2 were conserved in group II and were required for proper plastic localization of MSL2 (Jensen and Haswell 2012). Therefore, although the conserved pore-lining helices of MSLs are conserved, the important residues are different among different groups, indicating their different channel activities and functions.

**Figure 6.**
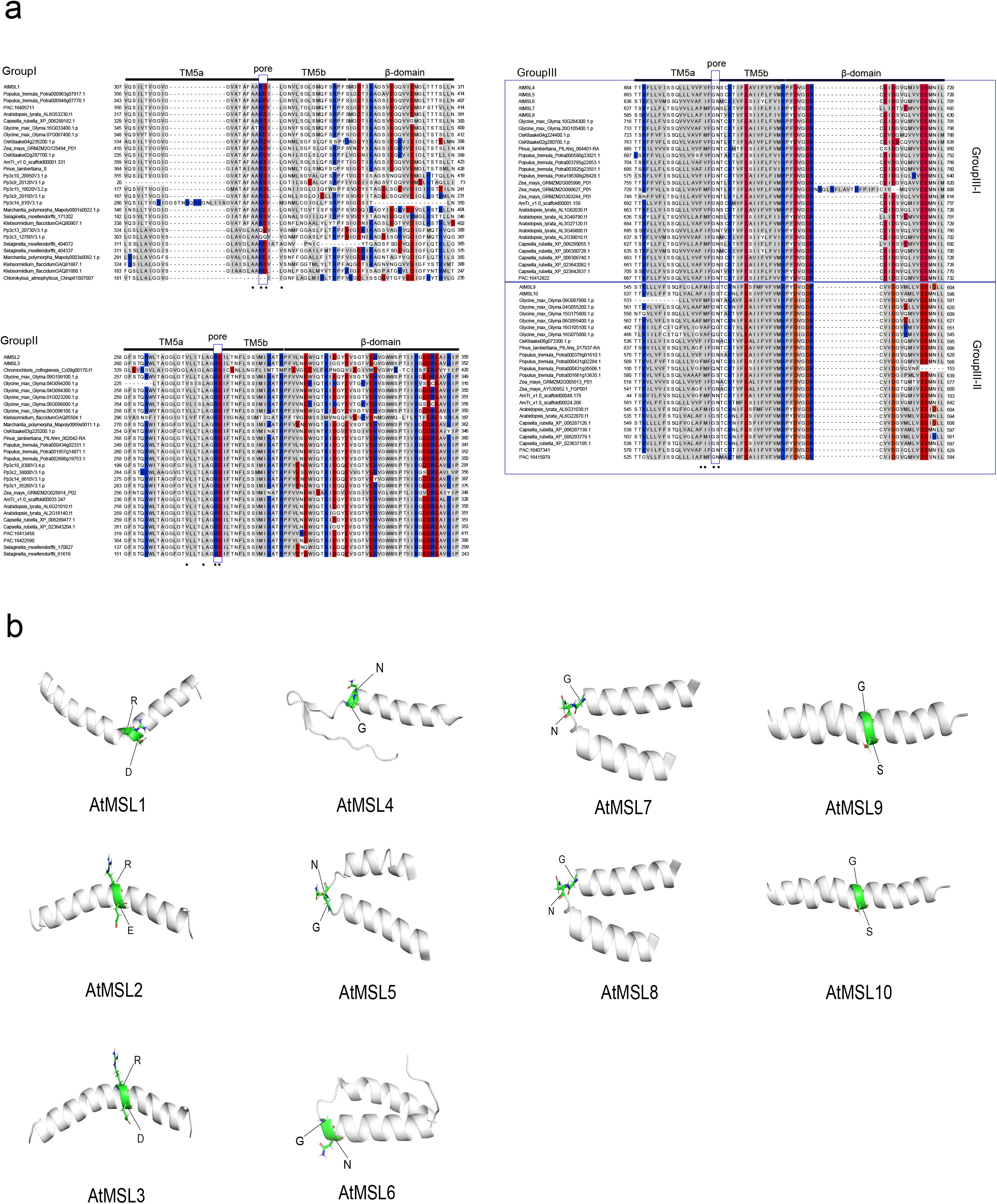
Multiple sequence alignment and conserved motifs in MSL proteins. (a) Alignment of pore-lining helices from MSL proteins. Sequences corresponding to TM3a, TM3b, and the ß-domain of *E*.*Coli* MscS are indicated by lines. Polar residues are white, non-polar residues gray, positively charged residues blue, negatively charged residues red. (b) Side view of the predicted pore-lining domain of the AtMSL1-10 created with I-TASSER. The side chains that predicted to form a kink are indicated with green sphere.

### Gene structure, conserved motif and phylogenetic analysis of MSLs in Brassicaceae

To explore the expansion of MSLs in Brassicaceae, we constructed a phylogenetic tree using Database III, which includes 10 Brassicaceae plants and 1 Brassicales plant (*Carica papaya*) (Figure 7). The phylogenetic tree showed that MSL4-8 and MSL9/10 were expanded in Brassicaceae, consisting with the former phylogenetic analysis (Figures 7). The number of exon/intron in Brassicaceae MSLs was analyzed (Figure 7). In Group I, most MSLs (4/7) have 5 introns. In Group II, most MSLs (12/15) have 12 introns. In Group III, more than two thirds of MSLs (51/64) contained 4 introns. Therefore, MSL members belong to different groups displayed different exon/intron structures, while MSL members belong to the same group showed similar exon/intron distribution (Table S2).

**Figure 7.**
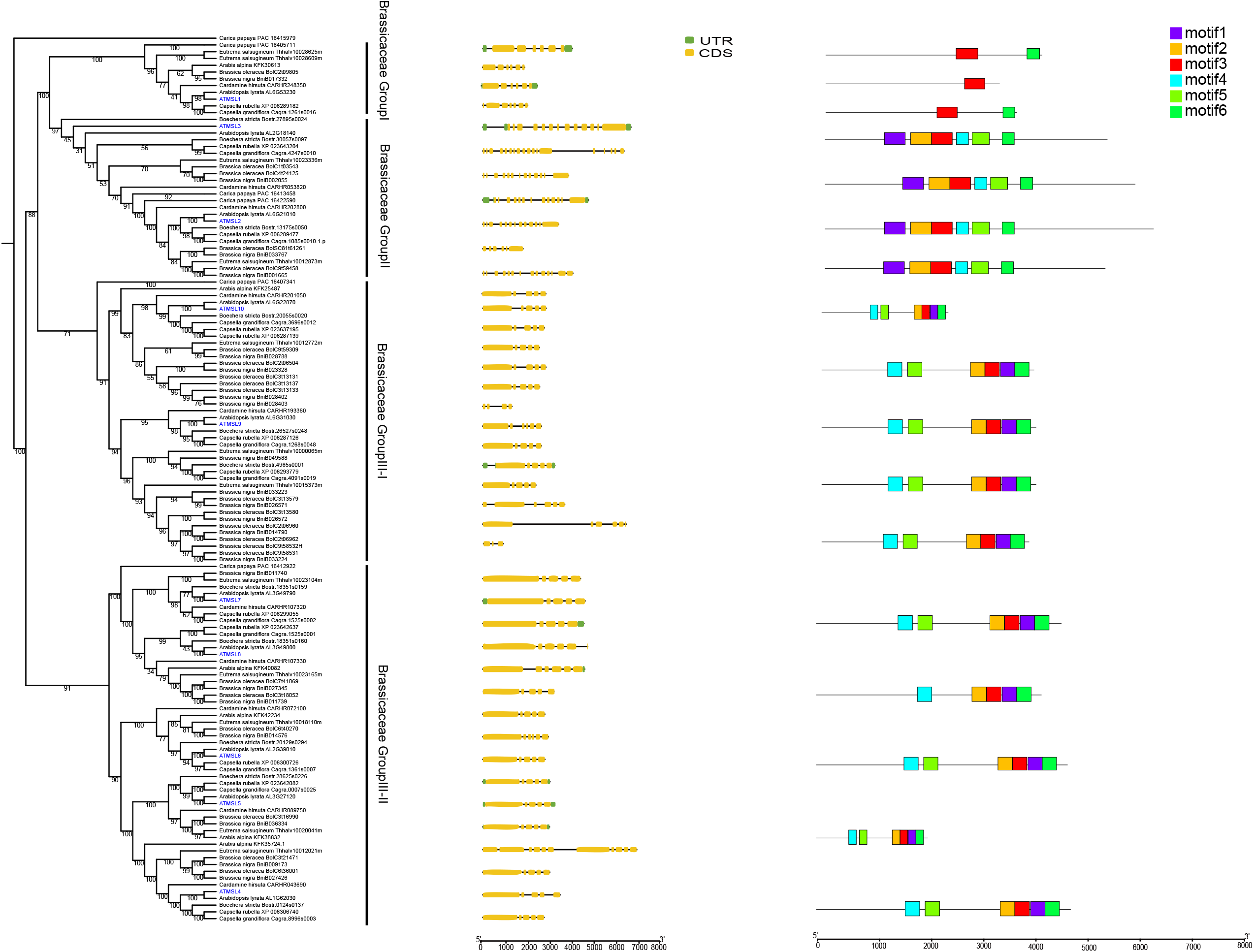
Gene structure, conserved motif and phylogenetic analysis of MSLs among Brassicaceae species. The phylogenetic tree was constructed using IQ-TREE with the parameter ‘ -m MFP -bb/alrt 1000 ‘ and 1000 ultra bootstrap replicates. The green boxes represent UTRs, yellow boxes represent CDSs and thin black lines represent introns. The motif in MSL proteins were identified by MEME program. Different motif numbered 1-6 has different colors.

The conserved motif was predicted by the MEME tool (Figure 7). MSL members belonging to Group I only have 1 (motif 3) or 2 motifs (motif 3, and motif 6). Most MSL members in Group II and Group III have 6 motifs. However, the motif locations are different between Group II and Group III MSLs. Both gene structure and protein motif results showed that MSL genes in the same group had similar gene structure and motif composition, indicating that there have similar functions. In addition, MSL genes between different groups display significant difference in both gene structure and motif composition, indicating the different functions.

### Expansion pattern of *MSLs* genes

Whole genome/segment and tandem duplications contribute significantly to the expansion of gene families, and gene duplication promotes genome evolution (Moore and Purugganan, 2003). To explore the expansion of *MSL* genes, we conducted a comprehensive synteny analysis (Figure 8). The results showed that the Arabidopsis *MSL* genes had 3 segmental duplication events (*AtMSL4/AtMSL5, AtMSL4/AtMSL7, AtMSL9/AtMSL10*), and one tandem duplication event (*AtMSL7/AtMSL8*). In addition, all these expanded *MSL* genes belongs to group III of MSL (Figure 8). Similar expansion patterns were also identified in other Brassicaceae plants, including *Arabidopsis lyrate, Brassica niqra*, and *Brassica oleracea* (Figure 8).

**Figure 8.**
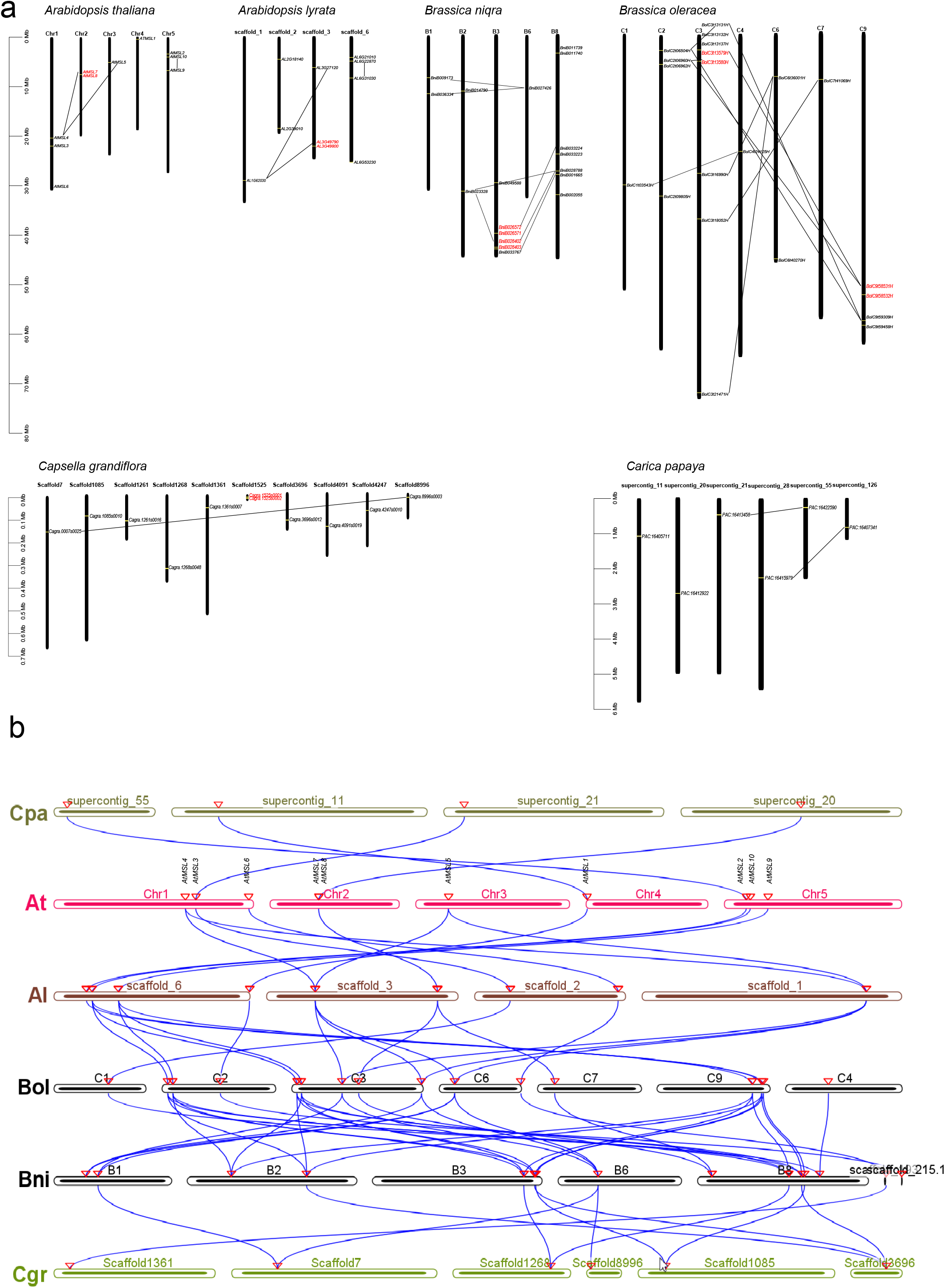
Synteny analysis of Brassicaceae MSL Proteins. (a) Genomic distribution of *MSL* genes across Brassicaceae species. Collinearity genes are linked by black line and tandem genes are marked by red color. (b) Multicollinearity analysis of *MSL* genes among different Brassicales species. The blue lines represent the MSL syntenic genes among *Carica papaya* (*Cpa*), *Arabidopsis thaliana* (*At*), *Arabidopsis lyrate* (*Al*),*Brassica niqra* (*Bni*), *Brassica oleracea* (*Bol*), and *Capsella grandiflora* (*Cgr*) genomes. The location of the *MSL* gene on the chromosome is marked by a triangle.

Furthermore, we performed a multicollinearity analysis of *MSL* genes among 6 Brassicales species (*Carica papaya, Arabidopsis thaliana, Arabidopsis lyrate, Brassica niqra, Brassica oleracea*, and *Capsella grandiflora*) to reveal the robust orthologs of these *MSLs* (Figure 8). The results showed that among Brassicaceae species, all *MSL* genes were collinearity. In addition, a relatively low collinearity was found between *Carica papaya* and other Brassicaceae species. These results indicate that the *MSL* genes were conserved and have the same ancestors.

## Conclusion

In this study, we performed comprehensive evolutionary analyses of *MSL* gene family in green plants. The phylogenetic insight provides valuable information for future molecular and biological investigations of various MSL proteins.

Our search show that the MSL proteins is widely exist in green lineage of plants and the copy number of MSL is varies among different species. Three MSL proteins were identified and divided into three different clades in chlorophyte, suggesting that MSLs may have diversified to possess complex localizations and functions in unicellular algae as in higher plants. During evolution, the MSL family expanded and formed four clades in seed plants. Based on the comprehensive analysis of MSL proteins among plant species, we propose an evolutionary model for MSL in green plants (Figure 9). Three ancestors of MSL exist in early hydrobiontic alage. Subsequently, these three MSL clades evolved independently during the evolution in land plants. The expansion of the MSL proteins in clade III occurred in seed plants, leading to two major branches of MSL4-8 and MSL9/10. However, the expansion of the MSL proteins in clade II occurred in dicots, leading to two major branches of MSL2 and MSL3. The divergence of MSL2/3 occurred after the monocot-dicot plant split; the divergence of MSL4/5/6/7/8 and MSL9/10 was the latest, before the emergence of Brassicaceae and after the Cleomaceae-Brassicaceae split. Within the MLS4-8 and MSL9/10 clades, a large expansion occurred in dicots, resulting in the formation of 2 and 3 subclades, respectively. This work provides insights that guide future investigations of MSL function in model and non-model organisms.

**Figure 9.**
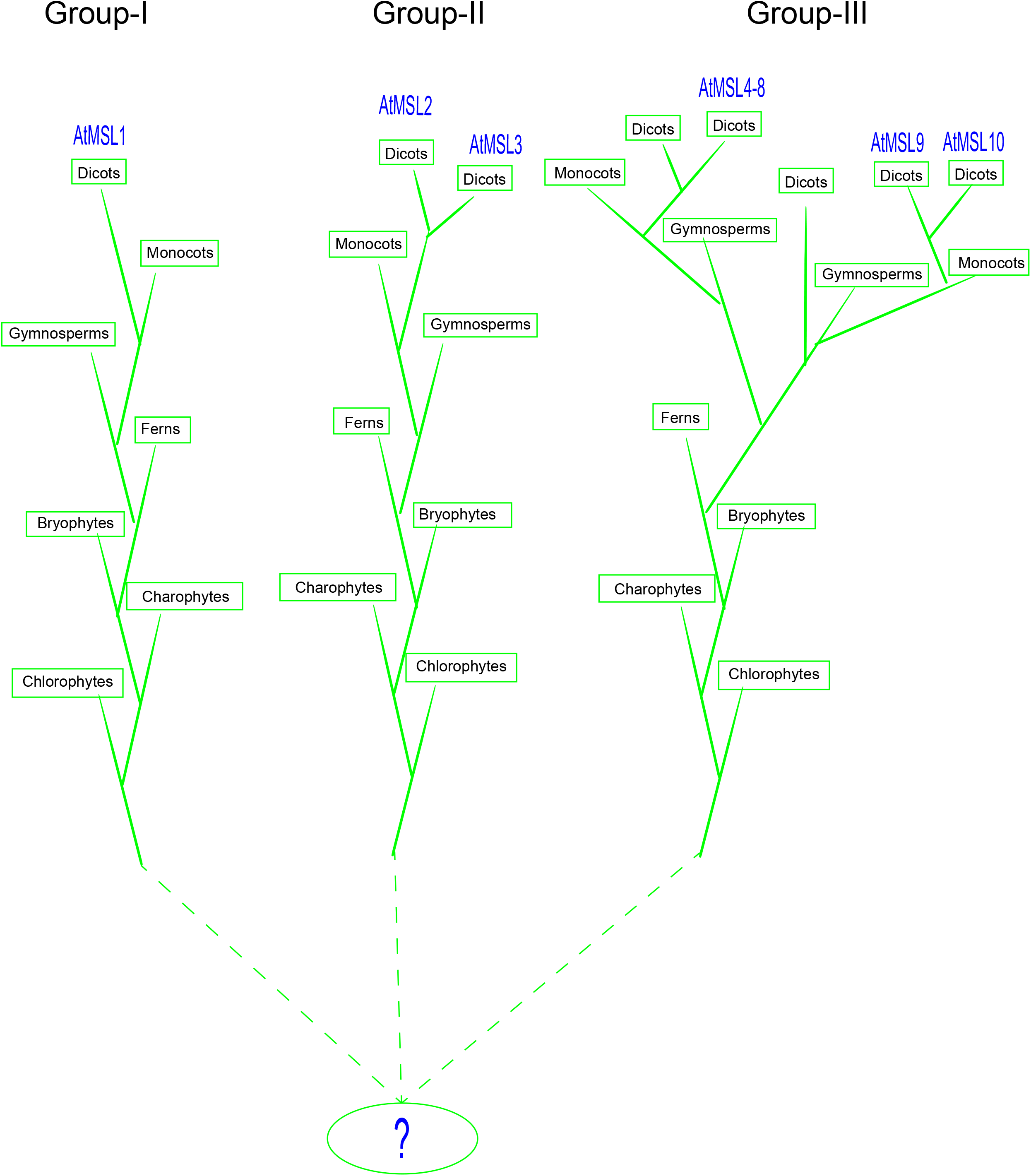
A proposed evolutionary model of MSLs in plants. The model is based on the phylogeny of MSLs and the cladogram of green plant species. The origin of plant MSLs was traced back to chlorophytes. During the evolution, MSLs were diverged into four major groups in angiosperms.

## Methods

### Data sources and sequence acquisition

A total of 176 species were selected and the corresponding genome sequences were obtained from public databases including NCBI (https://www.ncbi.nlm.nih.gov/), Phytozome v13. 0 (https://phytozome.jgi.doe.gov) and Gigadb (www.gigadb.org).

Two methods were used to identify *MSL* genes in plants. First, HMMER search (E-value=1e−10) was employed with the Hidden Markov Model profile of MscS channel domain (PF00924) to search the local Databases. Second, the amino acid sequences of *Escherichia coli* MscS and *Arabidopsis thaliana* MSL members were used to run a Basic Local Alignment Search Tool algorithms (BLASTP) search against the protein database with an E-value less than 10^−6^. The putative MSLs were further validated with online tools CDD (https://www.ncbi.nlm.nih.gov/Structure/cdd/wrpsb.cgi/) (Marchler-Bauer et al. 2015), HMM (https://hmmer.org/) (Finn et al. 2016) and SMART (https://smart.embl-heidelberg.de/) (Letunic et al. 2012). In total, 2113 MSL proteins were identified and used for further analysis (Table S1).

### Multiple sequence alignment, protein structure predictions and phylogenetic analysis

Multiple sequence alignments of MSLs were performed with MAFFT software (Katoh and Standley 2013). The putative structures of AtMSL2-10 were predicted with I-TASSER prediction server using the AtMSL1 crystal structure as a template (Roy et al. 2010; Li et al. 2020). The crystal structure was visualized with PyMOL software.

To explore the phylogenetic relationship between different species, we constructed 3 phylogenetic trees based on taxonomy: Dataset I contains 7 non-angiosperms and 9 angiosperms, Dataset II contains 156 angiosperms, Dataset III contains 11 Brassicales species (Table S3). The phylogenetic tree was constructed based on the core amino acid regions corresponding to transmembrane domain 3 and the adjacent consensus sequence. The Bayesian tree was constructed with MrBayes 3.2.1 software with the fixed Whelan and Goldman model (Ronquist and Huelsenbeck 2003). The maximum likelihood (ML) phylogenetic tree was constructed with IQ-TREE with the parameter ‘-m MFP -bb/alrt 1000’ and 1000 ultra bootstrap replicates (Minh et al., 2020).

### Subcellular localization prediction, conserved motif and gene structure analysis

The subcellular localization of MSLs was predicted with Wolf PSORT (https://wolfpsort.hgc.jp). Conserved motifs were predicted with MEME (http://meme.nbcr.net/meme3/mme.html) (Bailey et al. 2009) and gene structures were displayed through Gene Structure Display Server (GSDS) (http://gsds.cbi.pku.edu.cn/index.php).

### Synteny analysis

The gene duplication landscape was obtained using the MCScanX with the default parameters (Wang et al. 2012) and the syntonic map was generated using CIRCOS with the putative duplicated genes were linked by the connection lines.

### Data availability statement

All raw sequencing data were downloaded from public database. The detailed information could be found in Supplementary Table 1.

## Acknowledgements

This work was supported by the grants from National Natural Science Foundation of China (32170351), and Nanhu Scholars Program for Young Scholars of XYNU. The authors declare that they have no competing interests.

## Authors’ contributions

ZB-Z designed the research. ZB-Z, and F-Y wrote the manuscript. F-Y, T-X, JH-C, JJ-C, YR-C, and T-L performed the identification of *MSL* genes, protein structure and evolution analysis. JS-L participated in manuscript preparation and revision.

## Additional files

**Supplementary Figure 1. Phylogenetic analysis of MSL proteins among fungi, protozoa, bacteria and plants**.

**Supplementary Figure 2. Phylogenetic relationship within the Group III of MSL proteins**.

**Supplementary Table 1. Green plants, sources of sequence and genome version information used in this study**.

**Supplementary Table 2. The number of exons/introns in Brassicaceae MSL proteins**.

**Supplementary Table 3. The detailed information of MSL proteins identified in the present study**.

